# The central renin angiotensin II system – a genetic pathway, functional decoding and selective target engagement characterization in humans

**DOI:** 10.1101/2023.03.20.533428

**Authors:** Ting Xu, Zhiyi Chen, Xinqi Zhou, Lan Wang, Feng Zhou, Dezhong Yao, Bo Zhou, Benjamin Becker

## Abstract

The brain renin angiotensin II system plays a pivotal role in cognition and neuropathology via the central angiotensin II type 1 receptor (AT1R), yet the lack of a biologically informed framework currently impedes translational and therapeutic progress. We combined imaging transcriptomic and meta-analyses with pharmaco-resting state fMRI employing a selective AT1R antagonist in a discovery-replication design (n=132 individuals). The AT1R was densely expressed in subcortical systems engaged in reward, motivation, stress, and memory. Pharmacological target engagement suppressed spontaneous neural activity in subcortical systems with high AT1R expression and enhanced functional network integration in cortico-basal ganglia-thalamo-cortical circuits. AT1R-regulation on functional network integration was further mediated by dopaminergic, opioid and corticotrophin-releasing hormone pathways. Overall, this work provides the first comprehensive characterization of the architecture and function of the brain renin angiotensin II system indicating that the central AT1R-mediates human cognition and behavior via regulating specific circuits and interacting with classical transmitter systems.

## Introduction

Since the discovery of the enzyme renin by Tigersted and Bergman in 1898, the architecture and functions of the peripheral renin-angiotensin system (RAS) have been extensively examined and the discoveries have been successfully translated into mechanistic treatments for somatic disorders.

Accumulating evidence suggests that the RAS also plays a pivotal role in the central regulation of cognitive and emotional functions, as well as in the neuropathology and treatment of neurological and mental disorders.^1–3^ However, an overarching framework to guide the functional characterization and therapeutic potential of the central RAS is currently lacking which in turn critically impedes translational research and treatment development.

The RAS has been traditionally considered as a hormone system that regulates peripheral systemic functions, in particular cardiovascular and renal homeostasis.^1^ The discovery of a local brain RAS - that is independent of the peripheral system - in the 1970ties kindled research on the role of the RAS in directly regulating brain processes.^3,4^ Over the subsequent years, accumulating evidence suggested an involvement of the RAS in affective and cognitive domains including stress, fear, motivation and memory,^5–8^ as well as in the pathology of stress-related and neurodegenerative disorders including Post-Traumatic Stress Disorders (PTSD), depression and dementia.^9–11^ The hormone angiotensin II (Ang II) and its actions at the angiotensin type 1 and type 2 receptor (AT1R, AT2R) critically mediate most functional effects, with the AT1R mediating both the peripheral and central actions of Ang II.^12^ The AT1R is densely expressed in many organs, including the heart, kidney and brain,^13–16^ while in the adult brain the expression of AT2R is comparably low. Selective and competitive AT1R blockers such as Losartan are widely used for the treatment of cardiovascular and renal diseases and have an excellent safety record.^17^

The central AT1R is a G-protein coupled receptor (GPCR) which exhibits a dense expression in subcortical, limbic and frontal brain regions,^1,3^and thus represents a promising candidate target for the regulation of cognitive and emotional processes. Animal models have discovered that the central AT1R regulates highly specific functions such as stress reactivity, motivation and threat avoidance.^18–20^Based on these animal models subsequent human pharmacological studies have demonstrated a role of the AT1R in reward and motivational processing,^7,8^emotional memory encoding and fear extinction learning.^6^The effects of the AT1R in these domains are likely mediated by its dense expression in regions involved in motivation, emotion and memory such as the thalamus, striatum, amygdala, hippocampus, and prefrontal cortex.^1,3^Human pharmacological fMRI studies support the functional relevance of these circuits such that the blood-brain barrier permeable selective AT1R antagonist losartan: 1) attenuated amygdala threat reactivity,^21^ 2) facilitated regulation of threat via amygdala-hippocampal and amygdala-medial prefrontal pathways,^6,22,23^and 3) enhanced ventral striatal prediction errors and coupling in the mesocorticolimbic circuits during motivational paradigms.^7,8^ Large population studies further indicate that (chronic) treatment with pharmacological AT1R blockers can reduce the incidence of PTSD symptoms.^10,24^ While these results underscore a highly promising therapeutic potential of the AT1R, a recent multi-center randomized controlled clinical trial failed to determine treatment effects of an AT1R antagonist on PTSD symptomatology.^25^This underscores the need for an overarching and biologically-informed overarching framework for the regulatory role of the central AT1R in human cognition and emotion to facilitate generation and testing of precise translational hypotheses.

Some of the complex neurofunctional effects of the AT1R may emerge in interaction with other neural signaling systems, with recent evidence suggesting that interactions with the corticotrophin releasing hormone, dopaminergic, opioid, and vasopressin systems mediating AT1R regulation in some emotional and cognitive domains. The AT1R in the hypothalamic paraventricular nucleus (PVN) and the striatum critically regulate vasopressin^26^ and dopamine (DA) release,^18,27^ and its pharmacological blockade enhances DRD1 expression but decreases DRD3 sensitivity.^28^ With respect to the functional interaction, it has been shown that mice with a deletion of the AT1R gene from corticotrophin releasing cells exhibited less freezing than wild-type mice during tests of conditioned fear expression, an effect that may be caused by a decrease in the consolidation of fear memory.^29^ Accumulating evidence moreover suggests that central angiotensin inhibition modulates reward processing, reciprocal social interaction and pain processing via interactions with the opioid system.^30,31^ The role of the AT1R in stimulating vasopressin^32^ release may additionally mediate effects on anxiety during unpredictable threats^33^ (for the anxiolytic potential of vasopressin in humans see Zhuang et al., 2021).^34^ Together these studies describe further biological pathways that may mediate the complex effects of the AT1R, however, how these systems interact with the central AT1R system to regulate complex human behaviors has not been systematically examined.

To guide the discovery of regulatory mechanisms of the AT1R on human cognition and behavior and to facilitate the utilization of the AT1R as a novel therapeutic target, we here developed and tested an overarching biologically informed framework. We employed a systematic multi-methodal characterization of the architecture and functional relevance of the AT1R by means of brain gene expression mapping, meta-analytic cognition characterization and a randomized parallel group placebo-controlled pharmacological resting state fMRI design including independent discovery and replication samples to facilitate a robust precision imaging approach. Specifically, we employed a data-driven transcriptomic and meta-analytic mapping approach to precisely characterize the brain gene expression of AT1R pathways in humans and an independent meta-analytical functional characterization. Next, we conducted a discovery-replication randomized placebo-controlled parallel group pharmaco-resting state fMRI trial (two independent samples with n=66 healthy individuals each) employing the selective AT1R antagonist losartan to identify neurofunctional target engagement of the AT1R on regional activation and network level communication. We further utilized a partial least squares (PLS) regression analysis to determine the link between AT1R-modulated brain functional connectivity (FC) variation and the corticotrophin-releasing hormone, dopaminergic, opioid, vasopressin systems. We hypothesized that: 1) the AT1R shows a dense gene expression in subcortical regions characterized by a role in emotion, memory and reward processes as determined by meta-analytic functional decoding; that pharmacological blockade of the AT1R via losartan, 2) modulates spontaneous neural activity in the identified regions with high receptor density, and 3) regulates the functional communication in meso-cortico-limbic networks via interactions with the dopaminergic and opioid system.

## Results

### Experimental design

We combined neuroimaging transcriptomic and a discovery-replication pharmacological resting state fMRI strategy in two independent samples utilizing a selective competitive AT1R antagonist (Losartan) to determine the distribution of AT1R pathways genes in the human brain, their functional relevance in terms of neurofunctional target engagement and potential interactions with other neurotransmitter systems (**Fig. 1**). To increase the robustness of our findings, two independent randomized placebo-controlled between-subject pharmacological resting state fMRI experiments were conducted and served as discovery and replication samples (discovery, n=66; replication, n=66). Results of the discovery sample are provided in the main text, while detailed replication results are provided in the **Supplement information**.

**Fig. 1.**
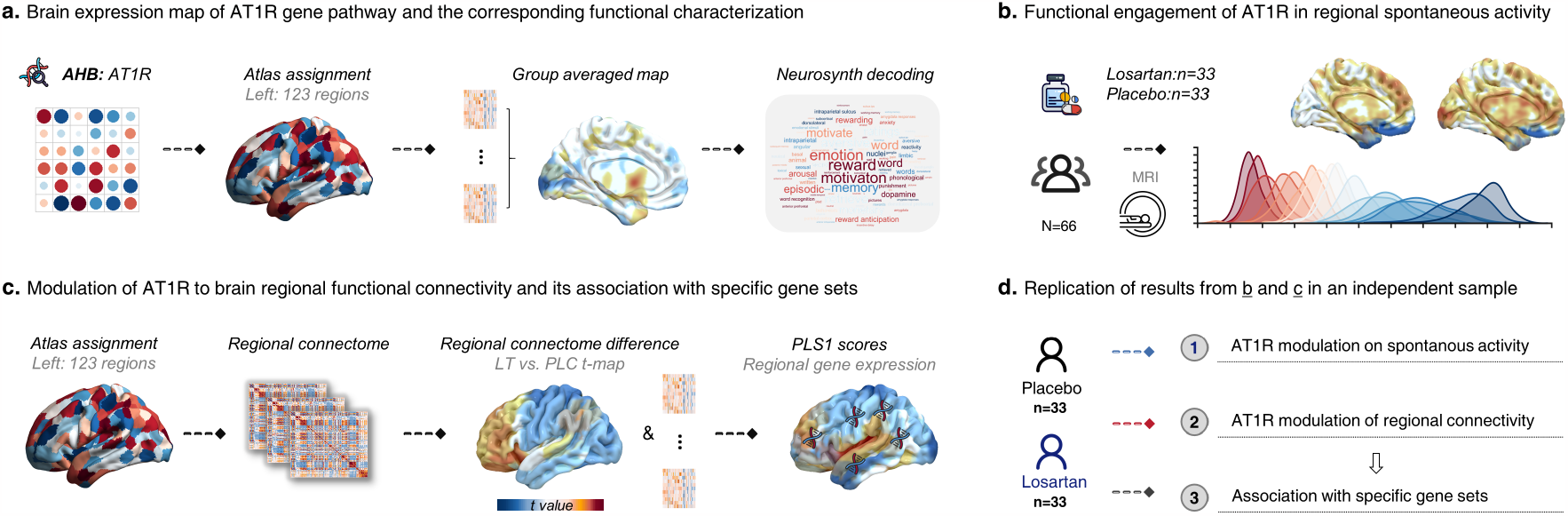
Schematic depiction of the analytic strategy. The study aims to: **(a)** characterize the architecture and function of the central AT1R pathways via transcriptomic maps from the Allen Human Brain Atlas from six postmortem brains in 123 brain parcels from the Brainnetome atlas (left hemisphere only in line with the gene expression profiles) and subsequent meta-analytic behavioral characterization of the resulting maps by means of the NeuroSynth database, **(b)** examine the molecular modulation of the AT1R pathways via a selective pharmacological blockade of the AT1R (versus placebo) with concomitant resting-state fMRI acquisition to determine target engagement of AT1R on spontaneous brain activity (restricted to left brain regions in line with the gene expression profiles), **(c)** explore interaction of the AT1R signaling pathways with other transmitter systems by means of functional connectivity and the partial least square (PLS) regression that examined whether the expression of selected genes from different neurotransmitter systems were related to network level effects of AT1R blockade. **(d)** Replicating the findings in panel b and c with an independent dataset. Abbreviations: AT1R-angiotensin type 1 receptor, LT-losartan, PLC-placebo.

### AT1R gene expression pathways in the brain and meta-analytic cognitive decoding

We initially mapped AT1R gene expression patterns in the human brain by capitalizing on AT1R gene expression maps from six donor brains collected from the Allen Human Brain (AHB) Atlas^35^ and extracting the regional specific expression values of the AT1R based on precise brain parcellations as provided in the Brainnetome atlas.^36^ In line with previous studies using the AHB database the analyses were restricted to left hemisphere^37,38^ to utilize a more number of donors. Compared to the average brain wide expression, the AT1R gene exhibited a particularly dense expression in subcortical and frontoparietal networks, yet a comparably low density in limbic and posterior to anterior cortical midline regions that represent core nodes of the default mode network (DMN). Statistically significant higher expressions were observed in the subcortical network (t=2.61, p<0.05, 95% confidence interval (CI), [0.00, 0.45], **Fig. 2a**) encompassing the thalamus, striatum, amygdala and hippocampus, while significantly lower AT1R gene expression was observed in the limbic network (t=-3.16, p<0.05, 95% CI, [-0.27, -0.03], **Fig. 2a**) that included the orbital gyrus (lateral and medial), superior temporal gyrus (medial and lateral), inferior temporal gyrus (intermediate ventral and rostral), fusiform gyrus and parahippocampal gyrus. Meta-analytic functional decoding of the AT1R mRNA expression maps revealed a strong overlap between the neurofunctional expression maps and memory and motivation/reward associated domains, including retrieval, motivational, monetary, anticipation and reward (**Fig. 2b**). In addition, to a lesser extend an overlap with negative emotion processing domains including terms such as aversive, anxiety, punishment or arousal was observed.

**Fig. 2.**
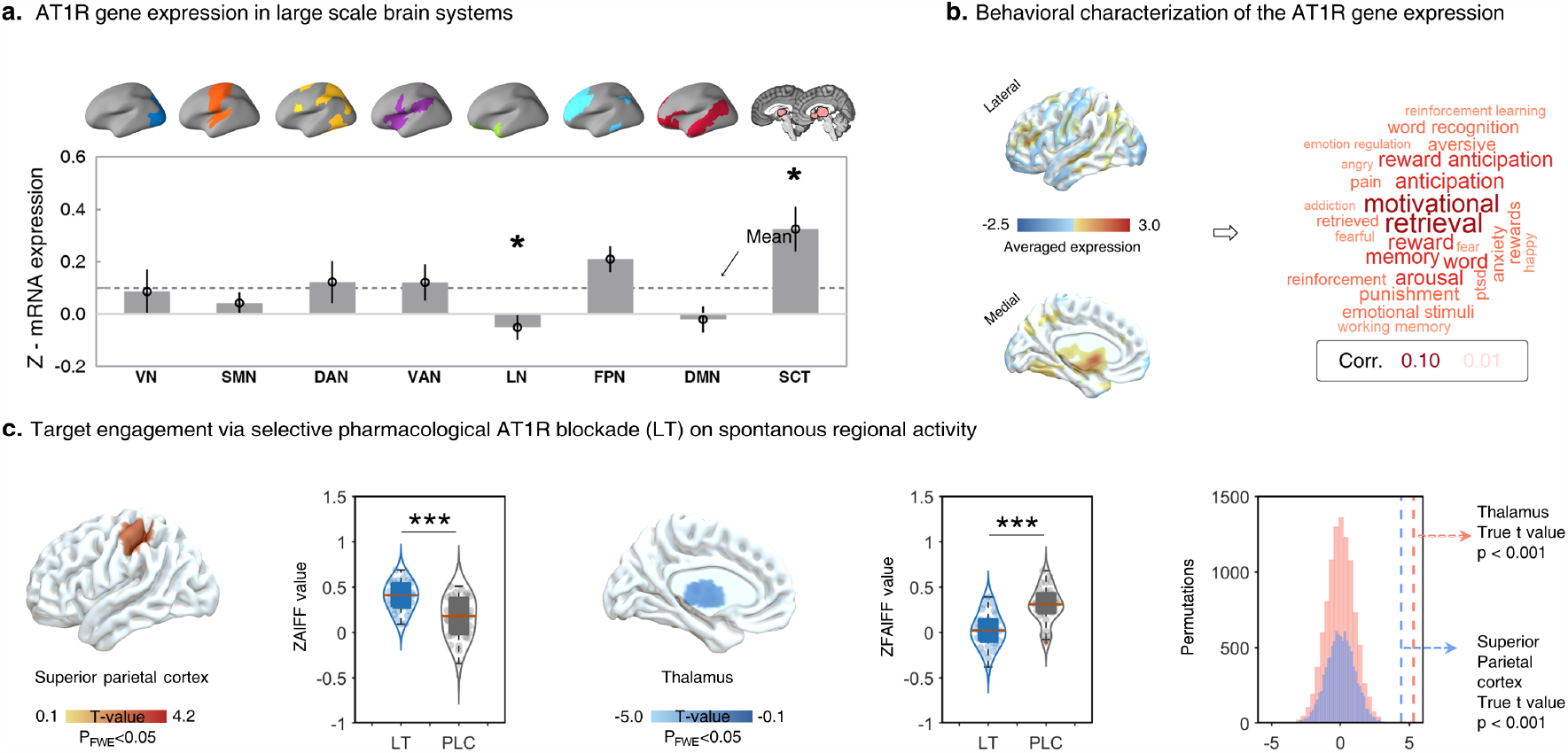
AT1R gene expression brain map and its meta-analytic functional characterization, as well as functional target engagement during pharmacological AT1R blockade on voxel-wise intrinsic brain activity. **(a)** Bars indicate the mean expression from the six donors with standard errors for a given brain network. The gray dashed line represents the mean expression across the entire brain. Relative to the average AT1R brain gene expression, the AT1R exhibited a higher expression in the subcortical (SCT)^*^ and fronto-parietal network (FPN) but a lower expression in the limbic (LN) ^*^ and default mode network (DMN). **(b)** Meta-analytic decoding of the central AT1R mRNA expression maps using the NeuroSynth meta-analytic repository revealed an association with functions in the memory and reward/motivation domains. **(c)** The AT1R antagonist losartan increased local intrinsic brain activity in the superior parietal cortex but decreased local spontaneous activity in thalamic regions characterized by a high expression of AT1R gene. ^***^indicates permutation test significance < 0.001. Abbreviations: AT1R-angiotensin type 1 receptor, LT-losartan, PLC-placebo, VN-visual network, SMN-somatomotor network, DAN-dorsal attention network, VAN-ventral attention network, LN-limbic network, FPN-fronto-parietal network, DMN-default mode network, SCT-subcortical network.

### Functional target engagement via a selective AT1R antagonist – effects on intrinsic activation and connectivity

To determine functional target engagement of the AT1R we conducted a randomized placebo-controlled between-group resting state pharmacological fMRI study using the selective competitive AT1R antagonist losartan. To precisely map the effects of acute AT1R blockade on the whole brain level with (voxel-level) spatial resolution treatment-induces changes in amplitude of low frequency fluctuations (fAlFF) were determined. fAlFF operates on the voxel-level and has been closely linked to spontaneous intrinsic neural activity and has a high sensitivity to metabolic activity, pharmacological receptor engagement and regional variations in gene receptor expressions.^39–42^ Comparing the losartan and placebo groups using voxel-wise two sample t-tests on zfAlFF of the left brain in the discovery cohort using a permutation test revealed that – compared to placebo – AT1R blockade significantly attenuated spontaneous activity in subcortical regions characterized by the highest AT1R gene expression, in particular the left mediodorsal (MD) and ventromedial (VM) thalamus (T=5.31, k=167, P_FWE-cluster_<0.05, peak Montreal Neurological institute (MNI): x,y,z= −6/−6/4, Fig. 3b). Losartan also increased spontaneous activity in the superior parietal cortex (T= 4.41, k= 448, P_FWE-cluster_<0.05, peak MNI: x,y,z= −34/−32/50, Fig. 3a), a region exhibiting a comparably low AT1R expression (see receptor maps presented in **Fig. 2b**).

**Fig. 3.**
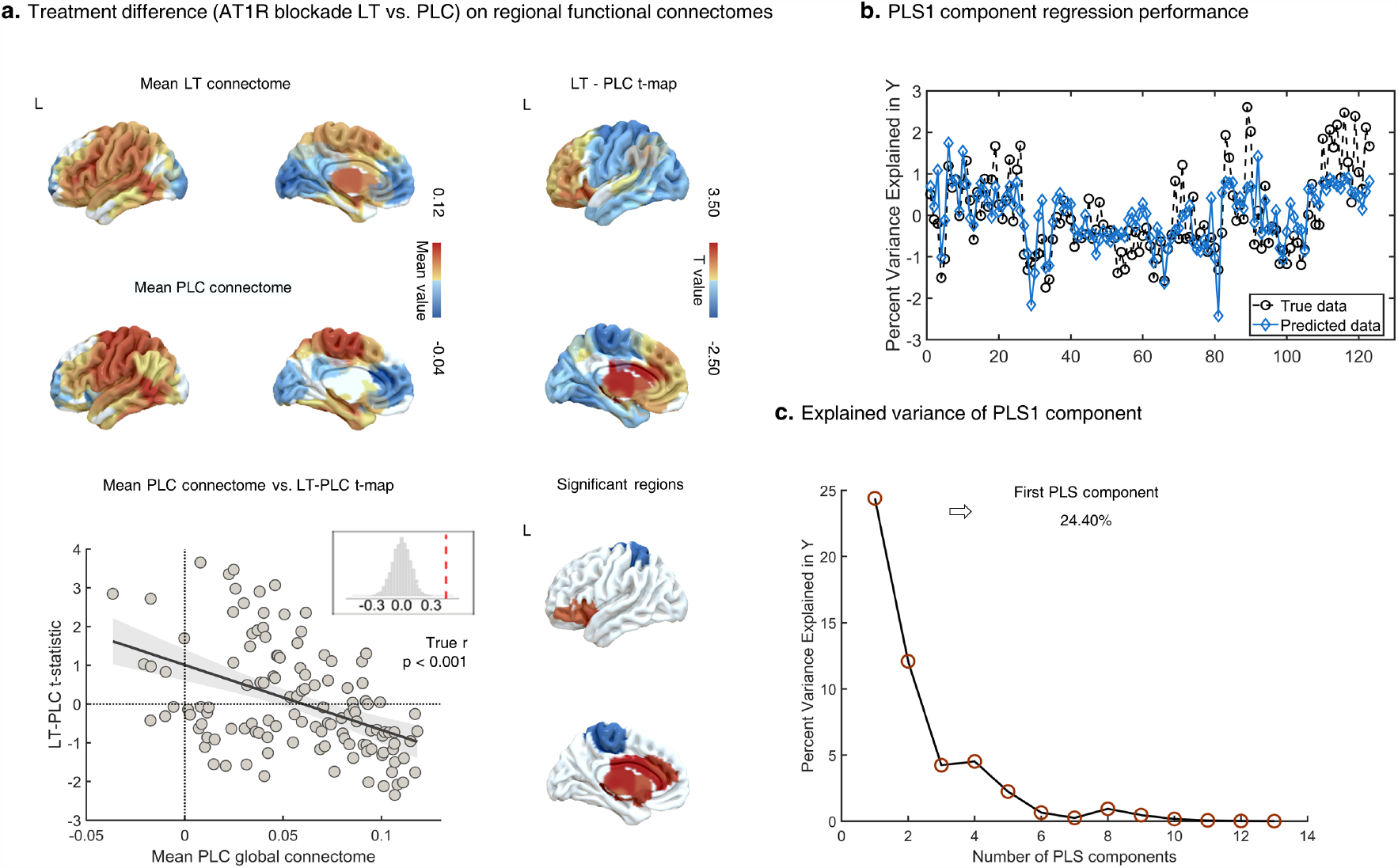
Effects of pharmacological AT1R blockade on regional functional connectivity. **(a)** Regional FC maps in the placebo and losartan groups, respectively, showing regional variations in FC on the whole-brain level, while the scatterplot shows the relationship between mean regional FC in the placebo group (x axis) vs. treatment effects as t statistic on FC (y axis). **(b)** Regression performance of first component from the partial least square (PLS) regression analysis. **(c)** Explained ratios for the 13 components obtained from the PLS regression analysis, but only the first component is selected due to the higher explained ratio (24.40%). Each component denotes a weighted linear combination of the expressions of the selected 13 genes. Abbreviations: LT-losartan, PLC-placebo, L-left.

We next determined the cortical maps of regional FC by coding the average distribution of regional FC in the placebo and losartan treated groups, respectively (**Fig. 3a**). Irrespective of treatment, FC in parietal and temporal regions was comparably high, while FC in frontal and occipital regions was relatively low. Mapping t statistics for the treatment differences for regional connectivity by cortical region (**Fig. 3a**) revealed a significant negative correlation between regional FC in the placebo group and significant treatment differences in regional FC (r= −0.45, p<0.001, **Fig. 3a**). This indicates that areas that normally exhibit the greatest regional FC (under placebo) exhibited the strongest decrease in FC following losartan, while conversely, regions with the lowest FC (under placebo) exhibited the strongest FC increase following losartan. Examining the pattern of effects on the regional level further revealed that losartan - compared to placebo - significantly increased FC in several key regions of the cortico-basal ganglia-thalamo-cortical circuitry (**Table S2, Fig. 3a**), including subcortical regions such as the MD and VM thalamus, caudate and putamen, as well as cortical regions such as the anterior cingulate cortex, medial and middle frontal gyrus, while it decreased FC in the superior parietal gyrus (**Fig. 3a**).

### Interaction of AT1R with other signaling systems and AT1R blockade-induced functional connectivity changes

Previous research indicates that the complex regulatory effects of the central AT1R system on cognitive and behavioral domains may evolve in interaction with other neural signaling systems and we thus focused on the corresponding gene expression maps:^9,29,31,43^ a dopaminergic set (DRD1, DRD2, DRD3, DRD4, DRD5), a corticotrophine release hormone set (CRH, CRHR1, CRHR2), an opioid set (OPRM1, OPRD1, OPRK1), and a vasopressin set (AVPR1A, AVPR1B). To explore functional interactions between ATR1 signaling and the other systems we employed a PLS regression approach to determine whether the expression patterns of the 13 selected receptor genes were related to the effects of AT1R blockade on regional FC. The first PLS component significantly explained 24.40% of the variance in the neurofunctional effects of acute AT1R blockade on regional FC (p<0.001, permutation test with 10,000 samples, **Fig. 3b**). The PLS1 gene expression weights were positively correlated with treatment differences on regional FC (r=0.49, p < 0.001, permutation test with 10,000 samples, **Fig. 4a**). The positive correlations indicate that genes positively weighted on the PLS1 component were densely expressed in regions exhibiting a marked FC increase after AT1R blockade losartan, while negatively weighted genes were densely expressed in regions exhibiting decreased FC after AT1R blockade (**Fig. 4b**). The DRD5 and OPRK1 coding dopaminergic and opioid systems, respectively, positively weighted on the PLS1 component associated with increased FC after AT1R blockade (**Fig. 4b**). The CRHR2 gene, an essential element in the physiological hypothalamic-adrenal stress response, negatively weighted on PLS1 linked with decreased FC after AT1R blockade (**Fig. 4b**).

**Fig. 4.**
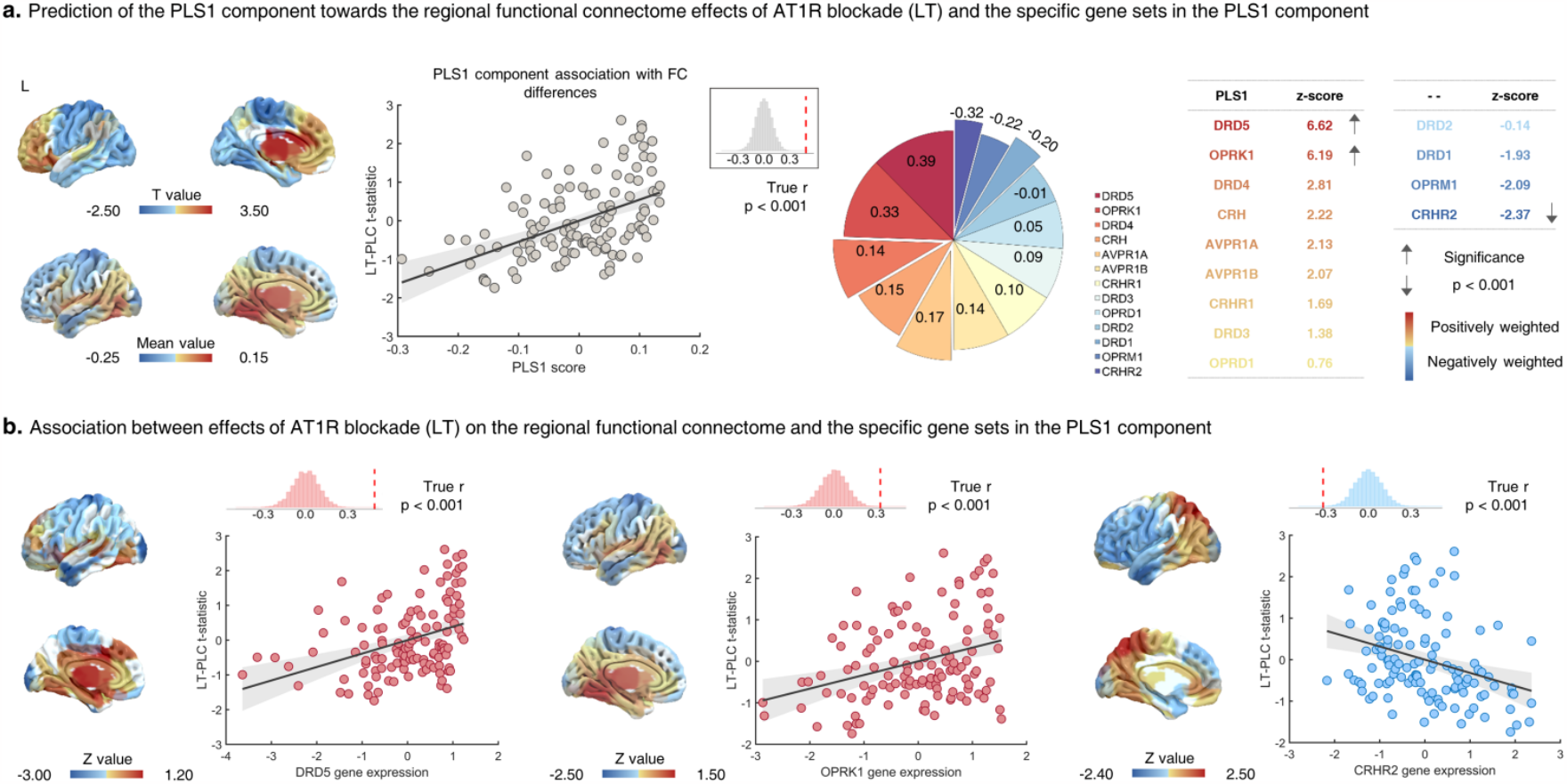
Determining interactions between ATR1 signaling with other neural signaling systems as mapped by gene sets and as a function of AT1R blockade induced neurofunctional effects on regional functional connectivity. **(a)** Cortical map AT1R-blockade induced changes in regional functional connectivity (FC) and the regional PLS1 scores (weighted sum of 13 gene expression scores). Scatterplot showing the association between regional PLS1 scores vs. treatment differences on regional FC. Genes with a strong positive weight on the PLS1 (i.e., DRD5, r= 0.39, p<0.001; OPRK1, r= 0.33, p<0.001) were correlated with increased FC, whereas genes negatively weighted on PLS1 (i.e., CRHR2, r = -0.32, p < 0.001) correlated decreased FC during acute AT1R blockade. **(b)** Gene expression maps significantly positively (i.e., DRD5, OPRK1) and negatively associated (CRHR2) with the neurofunctional effects of AT1R blockade on regional FC. Abbreviations: LT-losartan, PLC-placebo, L-left.

### Replication in an independent pharmaco-fMRI experiment

We also observed highly similar results to the observations in the discovery sample in the replication sample such that losartan-induced AT1R blockade specifically reduced intrinsic activity in the thalamus, but also modulated regional FC in subcortical-cortical pathways, while its associations with dopaminergic, opioid, corticotrophin gene sets have effects on regional FC were determined in line with findings in the discovery study (**Fig. S1-S3**).

## Discussion

The central renin angiotensin II system - in particular AT1R-mediated signaling - has received considerable interest as regulator of mental processes and highly promising treatment target for mental and neurological disorders.^2,25,44^However, the lack of an overarching biologically informed framework currently limits mechanistic and translational progress. Here we combined transcriptomic mapping with meta-analytic functional decoding and a randomized placebo controlled pharmacological resting state fMRI strategy in two independent samples to characterize the AT1R signaling pathways and their functional engagement and interaction with other transmitter systems comprehensively and systematically. Capitalizing on brain-wide transcriptomic gene expression data from the Allen Human Brain Atlas we mapped mRNA expressions specifically related to the AT1R and demonstrated that the receptor shows a particular dense expression in a subcortical network encompassing the thalamus, striatum and amygdalo-hippocampal formation. Meta-analytic behavioral decoding showed a strong association between the AT1R density distribution with memory and reward/motivational processes and to a lesser extend with negative affective processes. Pharmacologically induced acute blockade of the AT1R induced a suppression of spontaneous neural activity in regions characterized by a high AT1R expression, specifically the MD and VM thalamus, and a concomitant increase in spontaneous neural activity in the superior parietal cortex, a region that was characterized by comparably low AT1R expression. On the network communication level acute AT1R blockade increased functional connectivity between key nodes within cortico-basal ganglia-thalamo-cortical circuitry, specifically the MD and VM thalamus, caudate and putamen, as well as the anterior cingulate and adjacent medial prefrontal cortex. Applying PLS models to the AT1R functional engagement (blockade) and gene expressions maps of selected neural signaling systems demonstrated that the modulation of regional connectivity was associated with the topography of dopaminergic, CRH and opioid signaling systems, suggesting that some of the neural and behavioral functions of the AT1R emerge in interaction with these signaling systems. Replicating the analyses in an independent pharmacological fMRI study supported the robustness of the results in the discovery sample. Taken together, these findings provide the first comprehensive mapping of the central AT1R pathways and their functional relevance on the neural and behavioral level in humans.

### AT1R gene expression pathways in the human brain and their functional characterization

A precise mapping of the biological pathways of the central AT1R system in humans is lacking. Therefore, we first identified the brain topographical distribution of AT1R gene expression and found that the AT1R gene exhibits regional-specific variations in expression density, with subcortical regions encompassing the thalamus, striatum, hippocampus and the amygdala showing the highest expression of the receptor mRNA. Different lines of evidence from animal models suggests a (partly) convergent distribution of AT1R across species with dense expression in the striatum, hippocampus, amygdala and thalamus. The regional-specific AT1R expression may subserve the regulation of distinguishable behavioral domains such as: 1) AT1R G-protein-couple receptors in striatal glial cells affect inflammatory responses and in striatal projection neurons interact with dopamine neurotransmissions to modulate motor and reward-associated functions,^9^ 2) central AT1R blockade attenuates stress-reactivity of the hypothalamic-pituitary axis during isolation stress,^45^and deletion of AT1R from PVN of the hypothalamus attenuates anxiety-like behaviors,^46^ 3) AT1R eGFP+ expressed in the central amygdala gate the initiation of the fear response,^47^ and, 4) angiotensin II modulates hippocampal long-term potentiation (LTP) and associated memory processes via the AT1R.^48^ Consistent with the regional characterization of the AT1R distribution and function in rodent models the meta-analytic behavioral decoding of the AT1R distribution in humans revealed that the expression pattern aligned with brain systems involved in reward/motivation, memory, and negative affect.^49–52^ Previous studies have provided convergent evidence that the striatum plays an important role in reward and motivation and mediates the effects of pharmacological AT1R blockade in these domains.^7,8^ The hippocampus and amygdala have consistently been determined as brain systems critically involved in fear, anxiety, stress and emotional memory processes^53,54^ and mediate effects of AT1R modulation on fear and threat learning across species.^6,21-23,55,56^ The thalamus plays an important role in a range of sensory, cognitive and emotional processes including integration of sensory, emotional and conscious experiences and stress regulation and early angiotensin II studies in rodents indicate that this region may mediate a regulatory role of the RAS on somatosensory and stress processes.^57–59^ Together with the previous animal and translational models our findings converge on a regulatory role of the central AT1R systems in the domains of reward/motivation, memory, fear, and stress which may be mediated by neurofunctional effects in subcortical regions that exhibit a particular dense AT1R expression.

### Target engagement – effects of acute AT1R blockade on regional spontaneous activation and functional connectivity

To further map the neurofunctional engagement of the AT1R receptor we employed a resting state pharmacological imaging approach and observed that acute blockade of the AT1R induced regional-specific effects on spontaneous activity and regional connectivity in brain systems with a particularly high or low AT1R expression, respectively. We utilized losartan, a selective and competitive antagonist of the AT1R with the ability to cross the blood-brain barrier^60^ that following peripheral administration induced central neurofunctional effects in rodent models^61,62^ and following oral administration induced effects on task-related brain activity in humans.^6–8,21–23^The human studies revealed regional-specific neural effects that may mediate the effects of AT1Rs on specific task domains, however, the interaction between the task-specific neural engagement and the molecular action of the pharmacological agent limits conclusions with respect to overarching receptor engagement. Focusing on spontaneous activity and connectivity allowed us to map spatiotemporal engagement of the underlying systems irrespective of external demands.^63^ We found that acute pharmacological AT1R blockade suppressed spontaneous regional activity in subcortical regions with a high AT1R density, in particular the MD and VM thalamus, and concomitantly increased spontaneous activity in the superior parietal cortex, a region with comparably low AT1R expression density. Previous studies have demonstrated that the low-frequency fluctuation index examined in the present study is closely linked with spontaneous neuronal activity^64^and may bridge receptor expressions with neural activity.^42^ Animal models that examined effects of the AT1R on indices closely related to neural activity demonstrated that angiotensin II increases the discharge frequency within neurons of the lateral hypothalamus^65^and firing rates of hypothalamic PVN neurons^66^ – an effect eliminated by the AT1R antagonist losartan. Blockade of the densely expressed AT1R in the thalamus may thus reflect attenuation of excitatory effect of angiotensin II on thalamic neurons.

On the network level concordant but more widespread effects were observed such that pharmacological AT1R blockade enhanced functional communication in key nodes of the cortical-basal ganglia-thalamo-cortical circuitry and decreased connectivity in the superior parietal cortex. The FC measure applied reflects temporal associations between spatially distant neurophysiological events^67^and has been widely employed to determine functional relationship between neural regions.^68^Increased coupling within the cortico-basal ganglia-thalamo-cortical circuits may in turn reflect a higher temporal synchronicity between core rgeionss in these loops, while a decreased coupling within the superior parietal cortex may reflect decreasing synchronization within this system. The cortico-basal ganglia-thalamo-cortical loops represent a fundamental network organizational principle of the brain that critically mediates a broad range of functional domains ranging from motor and behavioral control towards learning and reward-based processes.^69–71^ Within this circuitry the MD and VM thalamus represent higher-order thalamic nuclei that relay and integrate an entire array of important signals,^72^ whereas the basal ganglia complex represents a key regulator of reward and motor related processes^73,74^ with bi-directional connections to the prefrontal cortex that facilitate adaptive control.^75^ Connections of the MD and VM thalamus with the basal ganglia and frontal cortex in particular support: 1) flexible and goal directed behavior including reward value and effort computations via thalamic-striatal connections,^76,77^ 2) regulatory control and decision making via thalamo-frontal circuits.^78,79^ Initial pharmaco-fMRI studies in humans furthermore indicate that an AT1R-blockade induced modulation of these circuits may mediate the regulatory role of the AT1R on reward, motivation and decision making in humans.^7,8^

### Network regulation of the AT1R and its association with dopaminergic, opioid and CRH systems

Given that several of the complex effects of the AT1R may be mediated by interactions with other transmitter systems we modelled the spatiotemporal overlap of neurofunctional AT1R effects and receptor maps of selected candidate signaling systems. Our findings indicated that specifically the DRD5, OPRK1 and CRHR2 gene expression showed a distinct association with AT1R effects on brain regional FC such that DRD5 and OPRK1 interacted positively with AT1R blockade induced network level changes, while the CRHR2 exhibited a negative association with the pharmacological blockade effects. Both brain dopaminergic and opioid systems contribute critically to reward and pleasure,^80,81^ while the CRH system is critically involved in mediating stress-related responses.^82,83^ Initial animal studies have begun to delineate the mechanisms underlying the interaction between the RAS and these neurotransmitter systems such that: 1) dopamine receptors and the AT1R show a regional specific high co-expression and regulate the release of dopamine in the striatum and hypothalamus^18,84^, 2) RAS and opioid interactions modulate the central regulation of peripheral nervous system actions of angiotensin II,^85^ 3) inhibition of central AT1R attenuates hypothalamic CRH-mediated isolation stress^86^and deletion of the AT1R gene from CR factor releasing cells attenuates conditioned fear expression.^29^ Together these findings underscore the interaction between the AT1R, and other neural signaling systems which may represent targets for modulating reward and stress domains. Impaired function of dopamine and opioid receptors and CRH-expressing neurons in subcortical regions ^87–91^ have been frequently associated with pathological reward processing deficits, and stress-related emotional dysregulations and may thus represent initial targets to translate AT1R modulation into clinical application.

### Limitations and outlook

Our findings need to be considered in the context of some limitations. First, the Allen human brain Atlas dataset provides a limited sample size and the donor group was variable with respect to age, gender and ethnicity. However, recent imaging transcriptomic studies indicate that this data is sufficiently generalizable^38,92^. Second, effects of neuropeptides are influenced by sex and trait-differences ^93,94^. While a recent study did not find sex-differential effects of losartan on neural activity^7^, future studies need to further determine potential effects of sex and trait variations. Third, while we capitalized on the robustness of a pharmaco-fMRI replication approach and consequently focused on convergent findings some differences were also observed.

## Conclusion

In summary, we provide the first biologically informed overarching framework for a comprehensive characterization of the biological and behavioral mechanisms of AT1R signaling across different methodological approaches. Across analyses the AT1R was densely expressed and modulated spontaneous neural activity and connectivity in subcortical-cortical circuits engaged in reward process, memory and fear/stress processing. Interactions with the dopaminergic, opioid and CRH systems may mediate some of the complex regulatory roles of the AT1R. The findings can inform future basic research and the translation of the AT1R as target for novel treatment approaches.

## Methods

### Participants

We employed a randomized parallel group placebo-controlled pharmacological resting state fMRI design including independent discovery and replication samples. N=70 healthy male volunteers were administered single oral doses of the selective competitive AT1R antagonist losartan (50mg) or placebo. Losartan crosses the blood brain barrier and its peak plasma levels are reached after 90 minutes while the terminal elimination half-live of losartan in brain activity and cognitive domains ranges between 1.5-2.5h^95,96^, which losartan effects on cardiovascular activity in heathy subjects were observed after 3h.^97^During the peak plasma period participants underwent resting state fMRI acquisition during which they were required to keep their eyes open. In each study an independent sample of seventy right-handed healthy male participants were enrolled. Given previous animal models and human studies reported sex differences in physiologic responses to RAS blockade^19,98^ and given that the vast majority of receptor maps for the AT1R in the Allen database were derived from male individuals,^35^ we decided to control for potential interactions between treatment and sex by only enrolling male participants in the present study.

All participants were free from current or a history of psychiatric, neurological, or medical disorders. Additional exclusion criteria included: excessive head movement (>2 mm translation or 2° rotation), current or regular use of psychotropic substances including nicotine, a body mass index <18 or >24.9, visual or motor impairments, and contraindications for MRI or losartan. A total of four participants were excluded due to excessive head movement (Losartan, n=2, Placebo, n=2) leading to a final sample of n=66 (mean±SD, age=20.94±2.44 years) included into the main analyses.

For the replication sample, N=70 healthy male participants were enrolled based on the same enrollment criteria. According to the enrollment and excluded criteria four subjects were excluded due to excessive head movement (Losartan, n=2, Placebo, n=2). Finally, the data of sixty-six subjects (mean±SD, age=20.90±2.06 years) were included into the replication analyses.

All participants provided written informed consent, and the study was approved by the local ethics committee at the University of Electronic Science and Technology of China (discovery sample) or Southwest University (replication sample). Study protocols adhered to the latest revision of the Declaration of Helsinki.

### Transcriptomic mapping - post-mortem brain samples

We capitalized on the full dataset of protein coding genes (n=20,737) from six donor brains as provided by the Allen Human Brain Atlas (http://human.brain-map.org/). Three donors were Caucasian males, one donor was a Hispanic female, and two donors were African-American males. Mean donor age was 42.5 (SD=11.2) years. Data was collected 22.3 (SD=4.5) hours after demise, on average (**Table S3 for donor profiles**). In line with the probe selected criteria reported in previous work^38,99^ we selected the probe with the highest differential stability which represented the probe with least amount of spatial variability among donors. The collection of human data complied with relevant ethical regulations, with institutional review board approval obtained at each tissue bank and repository that provided tissue samples. Moreover, informed consent was provided by each donor’s next-of-kin. Each donor’s brain was sampled in 363-946 distinct locations, either in the left hemisphere only (n=6), or over both hemispheres (n=2) using a custom Agilent 8×60K cDNA array chip. In line with previous works,^37,38^ the subsequent analyses were performed on left hemisphere to facilitate robust estimations.

### Creation of an AT1R gene expression brain map

The voxel-by-voxel volumetric expression map of the AT1R gene was created according to the procedure described in Quintana et al, 2019.^38^ To increase specificity sample locations and expression values of the AT1R gene were marked in native image space. Brain borders were subsequently mapped with the sample expression value that had the closet distance to a given order point. Next the space between scattered points into simplices was divided based on Delaunay triangulation, and then each simplex with values was linearly interpolated to yield a complete map. Finally, a composite brain map representing the average of 6 individuals for the left hemisphere was created. Next the resulting average brain was registered to the MNI stereotactic standard space using FSL linear registration and averaged so that each AT1R gene mRNA is represented in a single voxel-by-voxel brain map. For a more general characterization we additionally extracted averaged expression values for nine large-scale brain networks (i.e., default mode, fronto-parietal, somatomotor, limbic, dorsal attention, ventral attention, visual and subcortical networks) based on a total of 123 regions for the left brain that were based on the Brainnetome atlas^36^ (**Table S4**).

### Behavioral and cognitive characterization of the AT1R gene expression brain map

To determine the cognitive profile of the AT1R gene expression map, a reverse inference approach as implemented in the Neurosynth database^100^for meta-analytically decoding of functional domains was utilized. To this end the voxel-by-voxel mRNA map of AT1R was correlated with association Z maps, which reflects probability of a behavioral or cognitive process being present given knowledge of activation in a particular voxel.

### Potential confounders

In both discovery and replication sample, the losartan (Discovery, n=33; Replication, n=33) and placebo (Discovery, n=33; Replication, n=33) groups were comparable with respect to sociodemographics and cardiovascular indices arguing against nonspecific treatment effects (**Table 1 and Table S1**; all ps>0.05).

**Table 1.**
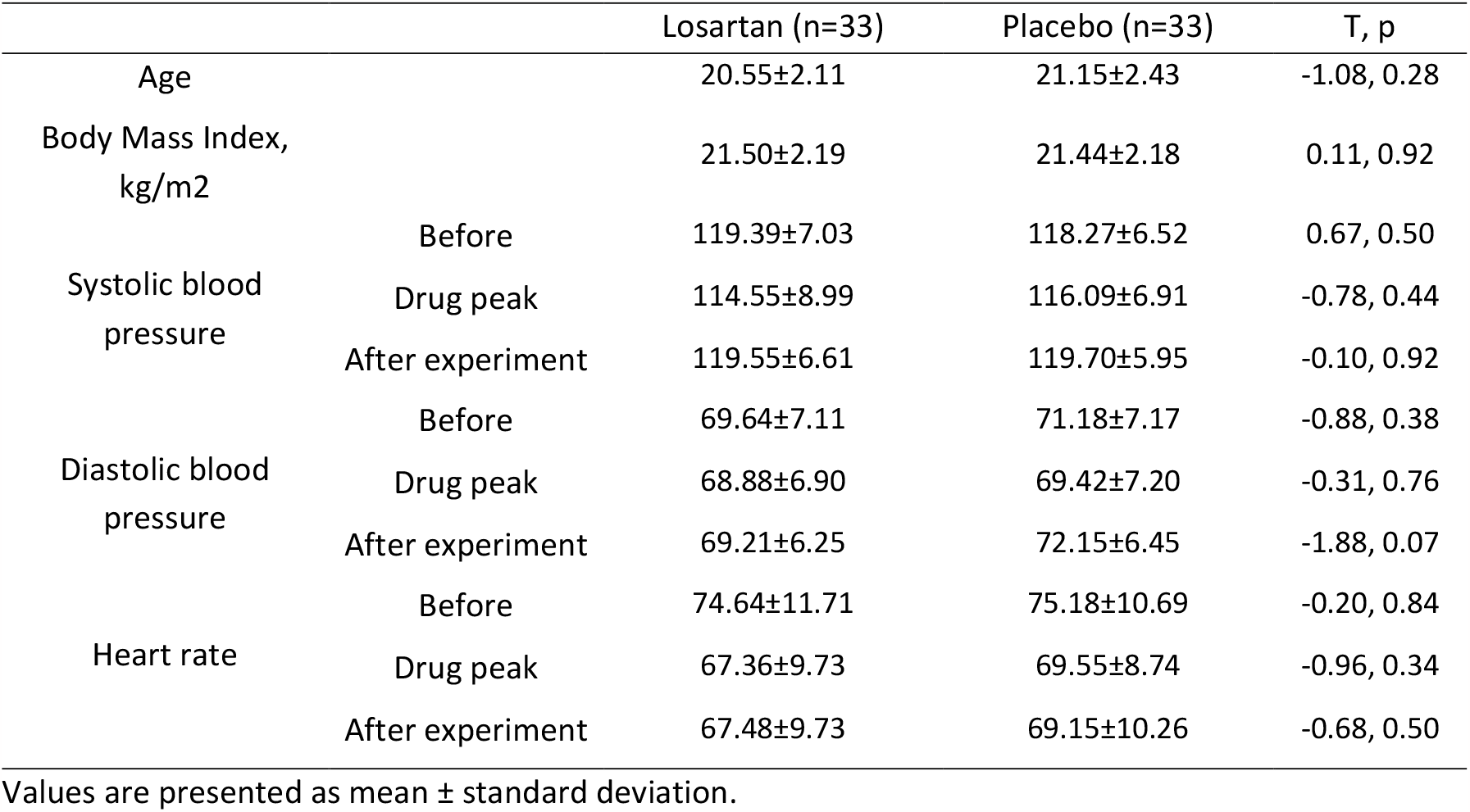
Demographics and potential confounders.

### MRI acquisition

MRI data of the discovery study were acquired on a 3.0-T GE Discovery MR system (General Electric Medical System, Milwaukee, WI, USA). The high-resolution brain anatomical MRI image were acquired using a T1-weighted sequence (repetition time=6ms; echo time=2ms; 156 slices; slice thickness=1mm; field of view=256×256mm; acquisition matrix=256×256; flip angle=12°; voxel size=1×1×1mm) to improve spatial normalization of the functional data and exclude subjects with apparent brain pathologies. Functional data using blood oxygenation level-dependent (BOLD) contrast were obtained on this scanner using a T2*-weighted echo planar imaging sequence (repetition time=2s, echo time=30ms; 39 slices; slice thickness=3mm; acquisition matrix=64×64; flip angle=90°; voxel size=3×3×3 mm).

For the replication study, the MRI data were obtained using a Siemens TRIO 3T system (Siemens, Erlangen, Germany). To improve spatial normalization and exclude participants with apparent brain pathologies a high-resolution T1-weighted image was acquired using a magnetization-prepared rapid gradient echo (MPRAGE) sequence (repetition time=1900ms; echo time=2.52ms; 176 slices; slice thickness=1mm; field of view=256×256mm; acquisition matrix=256×256; flip angle = 9°; voxel size=1×1×1mm). Functional MRI data was acquired using a T2*-weighted echo-planar imaging (EPI) pulse sequence (repetition time=2s; echo time=30ms; 33 slices; slice thickness=3mm; acquisition matrix=64×64; flip angle=90°; voxel size=3×3×3.6mm).

### MRI preprocessing

All MRI data were preprocessed using standardized workflows in fMRIPrep 20.2.1,^101^which is based on Nipype 1.5.1.^102^fMRIPrep is an automated pre-processing pipeline that flexibly employs tools from a variety of neuroimaging software packages. Many internal operations of fMRIPrep using Nilearn,^103^mostly within the functional processing workflow. For more details about preprocessing using fMRIPrep, please see **Supplement information**.

### Effects of ATR1 blockade on spontaneous neural activity

Previous studies demonstrated that low-frequency fluctuations within the 0.01-0.08 Hz frequency range capture spontaneous neuronal activity.^104,105^ We therefore created subject-specific fractional amplitude of low-frequency fluctuations (fALFF) maps via calculating a ratio of the ALFF within the specific low frequency band (0.01-0.08 Hz) to the total blood-oxygen-level dependent fluctuation amplitude within the full frequency band.^105^All fALFF maps were built based on left hemisphere to be in accord with previous analyses and were z-transformed before the statistical analyses (zfALFF). We subsequently conducted voxel-wise analysis on the level of the whole (left) brain using statistical parametric mapping software (SPM12, Welcome Department of Imaging Neuroscience, London, UK). Specifically, we compared the voxel-wise amplitude of the zfALFF between the losartan and placebo treatment groups by employing a voxel-wise two sample t test. The results were presented using p<0.05 at a cluster-level family wise error (FWE) correction with an initial cluster defining threshold of p<0.001 uncorrected at the voxel level.

### Effects of ATR1 blockade on regional functional connectivity

We constructed the FC matrix for the left brain using the approach as implemented in the high-order functional connectivity (HOFC) in BrainNetClass.^106^Specifically we extracted the regional averaged time series from each of the 123 left brain regions-of-interest (ROI) defined by the Brainnetome atlas^36^and calculated the Pearson correlation coefficient to indicate the low-order FC (LOFC) profile of the given ROI. Then HOFC is calculated as the similarity of LOFC profiles between each pair of ROIs, and could provide complementary information on topographical or spatial FC properties relative to the conventional LOFC.^107,108^This approach results in 123×123 HOFC connectivity profile for each subject. To examine the treatment differences, we used a general linear model with the HOFC matrix as the dependent variable, treatment as the independent variable, and age, BMI, cardiovascular activity indices (i.e., blood pressure and heart rate) as the covariates. Results were reported at a statistical significance threshold of p<0.05.

### Gene expressions related to the functional connectivity differences induced by the AT1R blockade

In line with our a priori hypothesis, we selected sets of mRNA of specific neural transmission systems that were reported to co-express and functionally interact with AT1R pathways to regulate neural transmission as well as behaviors including reward leaning, emotion and memory processes.^27,29,31,44^These sets included: a dopaminergic set (DRD1, DRD2, DRD3, DRD4, DRD5), a vasopressin set (AVPR1A, AVPR1B), a corticotrophin-releasing hormone set (CRH, CRHR1, CRHR2) and an opioid set (OPRM1, OPRD1, and OPRK1). We next utilized PLS regression analysis to determine associations between treatment-induced effects on FC differences (*t* statistics from the 123 cortical regions in the left hemisphere calculated based on the HOFC matrix comparing the losartan and placebo groups) and the gene expression for the selected 13 genes.

### Replication analyses

To replicate the results obtained in the discovery study, we repeated all analyses on resting-state MRI data, FC, and the transcriptomics profiles in the replication sample. Detailed results of the replication analyses are provided in the **Supplement information**.

## Data availability

Raw mRNA expression data is available from Allen Human Brain Atlas (http://human.brain-map.org/). The brain resting state data for some statistical analyses is available at https://figshare.com/projects/AT1R_brain_gene_expression/162517. The statistical figures were plot using Python (https://www.python.org/downloads/), Matlab 2020a (The MathWorks, Inc., Natick, MA; https://www.mathworks.com/) and R (https://www.r-project.org/) programming softwares. The brain maps were presented using python-based toolbox SurfIce (https://www.nitrc.org/projects/surfice/) and Matlab-based toolbox Canlabcore (https://github.com/canlab/CanlabCore).

## ACKNOWLEDGEMENTS

This work was supported by the China MOST2030 Brain Project (Grant No. 2022ZD0208500), the National Natural Science Foundation of China (Grants No. 32250610208, 82271583), and National Key Research and Development Program of China (Grant No. 2018YFA0701400). The authors express their gratitude to the Allen Institute for Brain Science for providing the gene expression data.

## AUTHOR CONTRIBUTIONS

This study was designed by TX and BB. All data were collected by TX, LW and XZ with the help of DY and BZ; TX, ZC and FZ performed data analysis and results integration with frequent discussions with BB. The manuscript was written by TX and BB and critically revised by all authors.

