## Supplements for "The central renin angiotensin II system – a genetic pathway, functional decoding and selective target engagement characterization in humans"

Benjamin Becker

University of Electronic Science and Technology of China

Xiyuan Avenue 2006, 611731 Chengdu, China


### MRI preprocessing

All MRI data were preprocessed using standardized workflows in fMRIPrep 20.2.1,[1,2] which is based on Nipype 1.5.1.[3] fMRIPrep is an automated pre-processing pipeline that flexibly employs tools from a variety of neuroimaging software packages. Many internal operations of fMRIPrep use Nilearn,[4] mostly within the functional processing workflow.

Utilizing tools from Advanced Normalization Tools (ANTs 2.2.0),[5] T1w images were corrected for intensity non-uniformity using the N4BiasFieldCorrection algorithm[6] and skull-stripped via a template-based brain extraction procedure (using the OASIS30ANTS template). Spatial normalization of T1w images to MNI space was performed through nonlinear registration with ANTs by using brain-extracted versions of both the T1w images and the ICBM 152 Nonlinear Asymmetrical template version 2009c.[7] Brain tissue segmentation of cerebrospinal fluid (CSF), white matter and gray matter was performed on the brain-extracted T1w images using FSL's FAST (FSL 5.0.9).[8]

Each participant's functional data was slice time corrected with AFNI's 3dTshift[9] and motion corrected with FSL's MCFLIRT.[10] This was followed by co-registration to the participant's corresponding T1w image using boundary-based registration[11] with nine degrees of freedom, via FSL's FLIRT.[12] Motion correcting transformations, the BOLD-to-T1 transformation and T1-to-template (MNI) warp were concatenated and applied in a single step using antsApplyTransforms (ANTs) with Lanczos interpolation, resulting in the functional data normalized to MNI space. From the MNI normalized functional data, mean signals were calculated from voxels within the CSF and white matter separately for use as nuisance regressors. Spatial smoothing was applied using an isotropic Gaussian kernel of 6 mm FWHM. Following smoothing, ICA-based Automatic Removal of Motion Artifacts (ICA-AROMA)[13] was performed to remove motion-related noise from the data. Tools from the DPABI toolbox (http://rfmri.org/dpabi)[14] were used to further denoise the MNI normalized, smoothed and ICA-AROMA denoised outputs from fMRIPrep. A band pass filter (0.01–0.08 Hz) was applied to the functional data, preceded by linear detrending and nuisance regression of subject-specific measures: mean white matter signal, mean CSF signal and functional volumes that had been identified as non-steady state outliers by fMRIPrep.

**Functional target engagement via a selective AT1R antagonist – effects on intrinsic activation and functional connectivity, as well as interaction with other specific neurotransmitter system**

To increase the robustness of the findings in discovery sample, we replicated analysis about intrinsic activation, functional connectivity (FC) and its association with specific neurotransmitter systems with an independent replication sample. Irrespective of treatment, FC in parietal and temporal regions was comparably high, while FC in frontal and occipital regions was relatively low (**Fig. S2a**). Mapping t statistics for the treatment differences for regional connectivity by cortical region (**Fig. S2c**) revealed a significant negative correlation between regional FC in the placebo group and significant treatment differences in regional FC ($r= -0.53$, $p<0.001$, **Fig. 2c**). This indicates that areas that normally exhibit the greatest regional FC (under placebo) exhibited the strongest decrease in FC following losartan, while conversely, regions with the lowest FC (under placebo) exhibited the strongest FC increase following losartan. Examining the pattern of effects on the regional level further revealed that losartan - compared to placebo – specifically but not significantly increased FC in several key regions of the cortico-basal ganglia-thalamo-cortical circuitry (**Fig. S2b**), including subcortical regions such as the MD and VM thalamus, caudate and putamen, as well as cortical regions such as the anterior cingulate cortex, medial and middle frontal gyrus, while it decreased FC in the superior parietal gyrus (**Fig. S2b**).

To explore functional interactions between ATR1 signaling and the other systems we employed a PLS regression approach to determine whether the expression patterns of the 13 selected receptor genes were related to the effects of AT1R blockade on regional FC. The first PLS component significantly explained 23.61% of the variance in the neurofunctional effects of acute AT1R blockade on regional FC ($p<0.001$, permutation test with 10,000 samples, **Fig. S3a**). The PLS1 gene expression weights were positively correlated with treatment differences on regional FC (r= 0.49, p<0.001, **Fig. S3b**). The positive correlations indicate that genes positively weighted on the PLS1 component were densely expressed in regions exhibiting a marked FC increase after AT1R blockade, while negatively weighted genes were densely expressed in regions exhibiting decreased FC after AT1R blockade (**Fig. S3c-d**). The DRD5 and OPRK1 coding dopaminergic and opioid systems, respectively, positively weighted on the PLS1 component associated with increased FC after AT1R blockade (**Fig. S3e**). The CRHR2 gene, an essential element in the physiological hypothalamic-adrenal stress response, negatively weighted on PLS1 linked with decreased FC after AT1R blockade (**Fig. 3e**).

**Table S1. Demographics and potential confounders in Discovery sample**

| | | Losartan (n=33) | Placebo (n=33) | T, p |
|---|---|---|---|---|
| Age | | 21.15±2.24 | 20.67±1.87 | 0.96,0.34 |
| Body Mass Index, kg/m2 | | 21.54±2.50 | 21.74±2.70 | -0.31,0.76 |
| Systolic blood pressure | Before | 111.39±7.81 | 111.42±4.19 | -0.02,0.98 |
| | Drug peak | 110.30±8.80 | 108.48±4.92 | 1.04,0.30 |
| | After experiment | 110.91±8.99 | 110.03±5.21 | 0.49,0.63 |
| Diastolic blood pressure | Before | 72.40±8.61 | 71.64±6.79 | 0.4,0.69 |
| | Drug peak | 68.79±8.05 | 70.58±7.17 | -0.42,0.68 |
| | After experiment | 70.76±7.72 | 71.94±7.03 | -0.65,0.52 |
| Heart rate | Before | 81.82±13.02 | 79.76±11.44 | 0.68,0.50 |
| | Drug peak | 76.00±11.08 | 74.21±10.81 | 0.66,0.51 |
| | After experiment | 74.67±10.58 | 74.82±10.41 | -0.06,0.95 |

Values are presented as mean ± standard deviation.

**Table S2. Brain regions of the thalamo-striato-cortical circuitry**

| Regions | x | y | z | Peak-voxels | T statistics |
|---|---|---|---|---|---|
| Caudate | -6 | 18 | -6 | 3885 | 3.65 |
| | -14 | -2 | 6 | | 3.47 |
| | -12 | -12 | 6 | | 3.47 |
| Anterior cingulate | -6 | 34 | 2 | 614 | 3.65 |

**Table S3. Detailed donor profiles**

| Ethnicity | Gender | Age | Post-mortem interval(hours) | Number of brain samples | hemisphere |
|---|---|---|---|---|---|
| Caucasian | Male | 57 | 25.5 | 363 | Left |
| Caucasian | Male | 31 | 17.5 | 529 | Left |
| Hispanic | Female | 49 | 30.0 | 470 | Left |
| Caucasian | Male | 55 | 18.0 | 501 | Left |
| African American | Male | 24 | 25.0 | 946 | Both |

| | | | | | |
|---|---|---|---|---|---|
| African American | Male | 39 | 18.0 | 893 | Both |

**Table S4. 123 brainnetome regions only for left brain hemisphere**

| Network | Regions |
|---|---|
| Default mode network | SFG,MFG,IFG,OrG,STG,MTG,ITG,PhG,pSTS,IPL,PCun,CG |
| Frontal parietal network | SFG,MFG,IFG,OrG,STG |
| Somatomotor network | SFG,PrG,PCL,STG,PoG,INS |
| Limbic network | OrG,STG,ITG,FuG,PhG, |
| Dorsal attention network | SFG,MFG,IFG,MTG,ITG,FuG,SPL,IPL |
| Ventral attention network | IFG,PrG,PCL,IPL,PCun,INS,CG |
| Visual network | FuG,PhG,MVOcC,LOcC |
| Subcortical network | Amyg,Hipp,BG,Tha, |

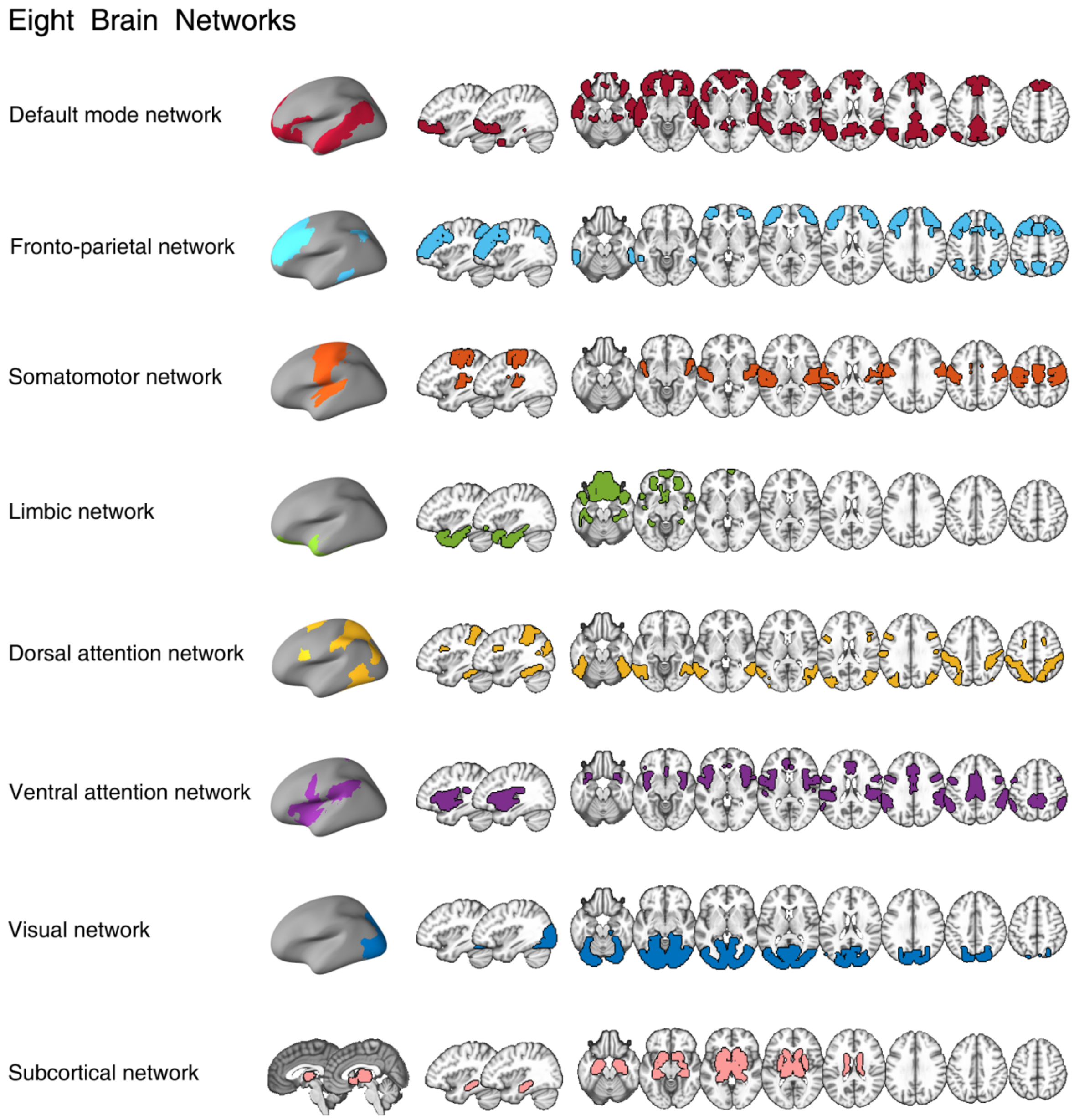


**Fig. S1** Brain networks generated from combination of Brainnetome atlas[16] with Yeo's Seven networks[19].

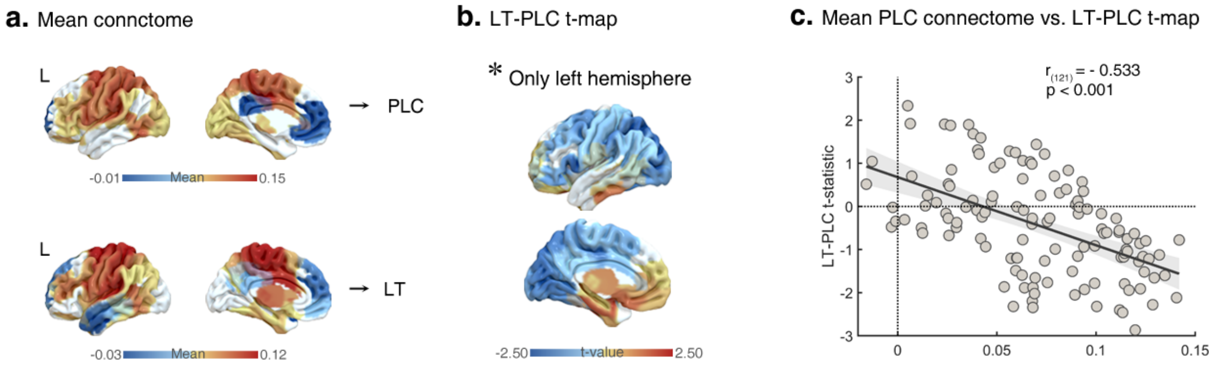

**Fig. S2** Effects of pharmacological AT1R blockade on regional functional connectivity in validated cohorts. **(a)** Regional FC averaged within the placebo and losartan groups, respectively. **(b)** Treatment effects in terms of t statistics for the treatment differences in FC across all brain regions. **(c)** The regional FC of placebo group (from b) is strongly negatively correlated with treatment differences on regional FC (r=-0.53, p<0.001). Abbreviations: LT-losartan, PLC-placebo,L-left.

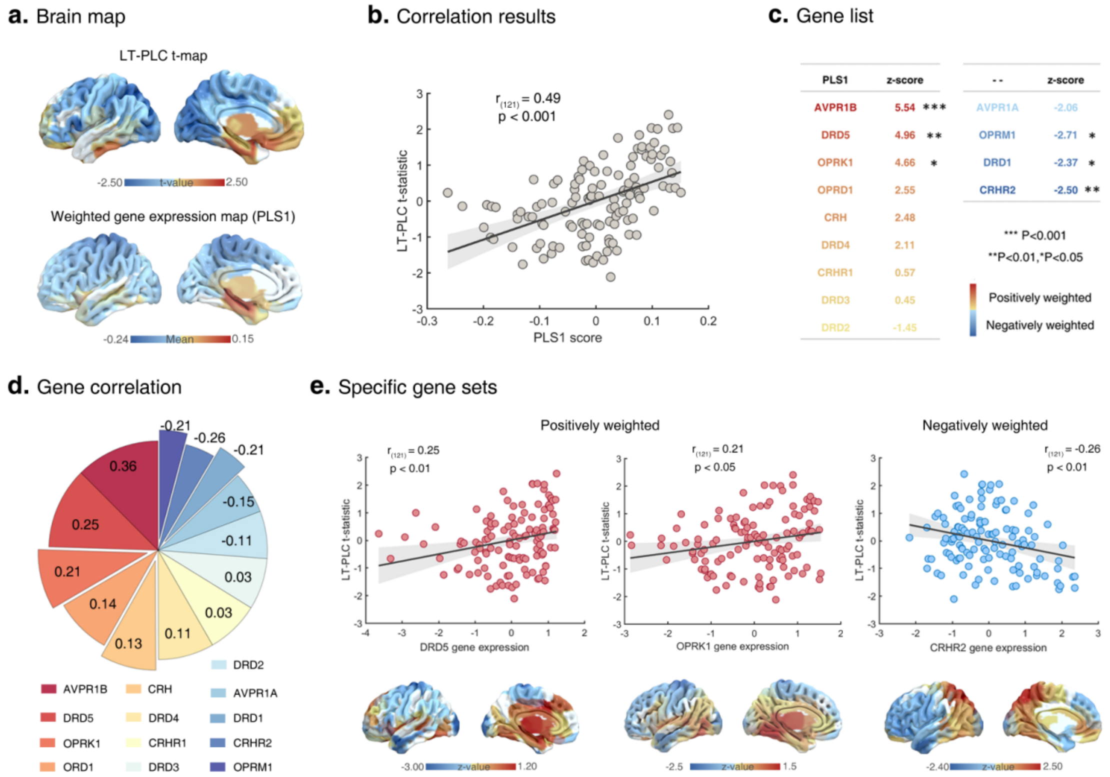


**Fig. S3** Determining interactions between ATR1 signaling with other neural signaling systems as mapped by gene sets and as a function of AT1R blockade induced neurofunctional effects on regional functional connectivity **(a)** Cortical map AT1R-blockade induced changes in regional functional connectivity (FC) and the regional PLS1 scores (weighted sum of 13 gene expression scores). **(b)** Scatterplot indicates the association between regional PLS1 scores vs. treatment differences on regional FC. **(c-d)** Genes with a strong positive weight on the PLS1 (i.e., DRD5, r=0.25, p<0.01; OPRK1, r=0.21, p<0.05) were correlated with increased FC, whereas genes negatively weighted on PLS1 (i.e., CRHR2, r=-0.36, p<0.01) correlated decreased FC during acute AT1R blockade. **(e)** Gene expression maps are significantly positively (i.e., DRD5, OPRK1) and negatively associated (CRHR2) with the neurofunctional effects of AT1R blockade on regional FC. Abbreviations: LT-losartan, PLC-placebo, L-left.
